# Enhancing information extraction from field potentials in electrophysiology studies

**DOI:** 10.1101/2024.11.18.624059

**Authors:** Shailaja Akella, Jose C. Principe

## Abstract

Multi-channel recordings from the brain serve as the primary approach to address various mechanistic and behavioral questions about neural activity and the underlying circuitry. While electrodes can detect neural activity at a distance from their source generators, activity from other concurrently active neural sources is also volume conducted to the electrodes, thus creating linear dependencies among the channels. These dependencies pose challenges in discerning specific communication patterns between different neuronal populations, and their behavioral implications. In this study, we demonstrate the capability of a marked point process (MPP) representation of channel activity, focusing on oscillatory bursts, to curtail the effects of volume conduction noise. By characterizing the localized spectral information within oscillatory bursts, we achieved a *∼* 45% reduction in channel correlations across three recording modalities of field potentials (electroencephalography, electrocorticography, and local field potentials). We further provide evidence that the implemented sparse representation preserves both behavioral and causal information in the signal. We illustrate our findings with two examples: 1) retention of finger-level movement information in field potentials recorded from humans, demonstrated using a simple online classifier, and 2) retention of top-down connectivity information between the prefrontal and motor cortices of behaving rats. Overall, our results underscore the novelty of using a marked point process representation of oscillatory bursts to concisely encode behavioral and connectivity information while attenuating the effects of volume conduction from the causal signal sources.

**Author summary:** Multi-channel recordings from the brain are critical to understanding how different brain regions communicate and drive behavior. However, the electrodes that detect neural activity also pick up signals from other active neural sources, leading to misleading linear dependencies among the channels. In our study, we present a novel approach using a marked point process (MPP) representation of channel activity using oscillatory bursts, to minimize the interference caused by volume conduction noise. By isolating the spectral characteristics within these oscillatory bursts, we achieved a *∼* 45% reduction in channel correlations across three different types of field potential recordings: electroencephalography (EEG), electrocorticography (ECoG), and local field potentials (LFPs). Importantly, our method not only reduces noise but also retains critical behavioral and causal information in the neural signals. Our findings reveal that a marked point process representation of oscillatory bursts provides a more precise and noise-resilient representation of neural activity. This advancement has significant implications for both understanding brain function and improving the accuracy of brain-machine interfaces.

## Introduction

Neuronal oscillations are emergent population-level field potentials that capture the temporal dynamics of the neuronal interactions at millisecond resolution. These oscillations are commonly monitored through non-invasive scalp electrodes (electroencephalography, EEG), invasive cortical surface electrodes (electrocorticography, ECoG), or electrodes implanted within the brain media (local field potentials, LFP). Regardless of the recording technique, neural activity detected at the electrodes results from the propagation of electric current through the conductive brain tissue between the generating dipoles and the measuring electrode. However, the finite impedance of the heterogeneous neural tissue causes the current flow to disperse through the brain media, such that the electrodes ultimately capture an attenuated, spatially averaged version of the source activity. This dispersion, known as volume conduction, introduces linear dependencies among channel activities at the electrode, leading to a loss of source specificity [1]. Addressing and mitigating these linear dependencies between channel activities is essential to accurately map the locations and interactions between various neural sources in the brain.

Consequently, a growing literature has focused on devising methods that suppress these confounding effects of volume conduction [1–4]. A prevalent approach, current source density (CSD), utilizes a surface Laplacian for spatial filtering [1]. However, its high susceptibility to noise, insensitivity to deep sources and requirement of a dense electrode array make it challenging to accurately estimate the spatial patterns of neural current sources. Capitalizing on the instantaneity of volume conduction effects, several researchers have introduced modifications on the existing coherence- and phase-based measures by excluding interactions that occur at a phase difference of 0° (or 180°) [5, 6]. While these measures offer insights into phase-lagged interactions, they may inadvertently filter out crucial connectivity information. Moreover, most coherence- and phase-based methods apply Fourier Transform techniques within finite observation windows, limiting their representation in both time and frequency space. Perhaps then a more refined strategy to mitigate the effects of volume conduction is to directly estimate source-level connectivity using blind source separation (BSS) methods [7].

These methods can approximate the anatomical location of the interfacing brain areas, however, unmixing sources is never perfect and often necessitates post-processing to distinguish important sources. Recognizing these challenges, there is a pressing need for methods that can effectively address the dynamic aspects of neural activity while ensuring an accurate interpretation of the observed neural signals.

In this paper, we introduce a novel representation-based technique aimed at addressing volume conduction by leveraging neurophysiological principles governing the generation of neuronal oscillations. Of particular interest is a recurring phenomenon immediately noticeable (by eyeballing) in the field potentials: appearance of a transient, distinctly organized, waxing and waning pattern that emerges from the featureless noisy background activity (Fig 1**A**). These brief oscillatory bursts are a direct consequence of synchronized synaptic interactions between the neural assemblies at the level of single neurons [8]. The significance of these phasic phenomena has been extensively recognized in the literature, serving as constructs of physiological and cognitive states, binding links for coherent perception, and influencing stimulus processing through top-down influences [9]. Moreover, many have argued for the role of bursts as a potential mechanism for cost-effective multiplexed communication between brain structures. Our approach rests on the premise that a rigorous characterization of these oscillatory bursts, in terms of their structure, sparsity, signal-to-noise ratio and transiency, will ultimately suppress volume conduction effects, where in general, the correlated activity will not adhere to their specific statistical properties. Insofar as, we replace the original heavily correlated single channel activity with a succinct marked point process (MPP) of oscillatory burst events.

**Fig 1.**
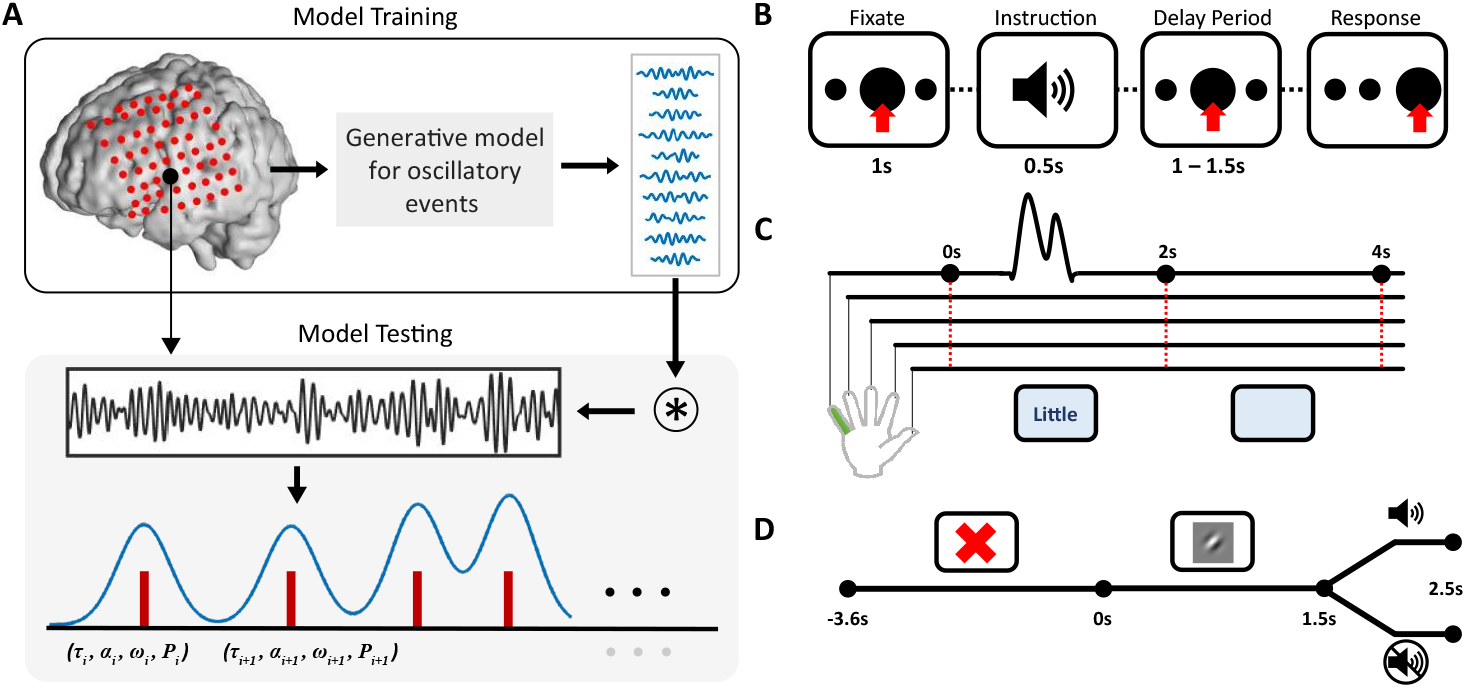
Transient model for oscillatory bursts. **A**, During the training phase (top), the generative model learns a dictionary of typical oscillatory patterns, *D* = *d*_*ω*_*ω* = 1^*K*^, from a subset of bandpassed LFP traces used as training data (top right panel). In the testing phase (bottom), the marked point process representation of a bandpassed LFP trace from the test set is constructed by convolving this trace with the learned dictionary, *D*. Once a burst is identified, its key properties or markers are identified as its time of occurrence (*τi*), amplitude (*α*_*i*_), power (*P*_*i*_), and the most representative dictionary template (*ω*_*i*_), allowing for a marked point process representation of oscillatory bursts (red). The M-spectrogram (blue trace) is constructed by weighting the intensity function of the point process by the transient power within the oscillatory burst at its time of occurrence. **B - D**,**Experimental setup for various recording modalities. B**, Dataset-1, Local field potentials were recorded from the prefrontal and motor areas of rats (n = 3, 10 sessions) performing a two-alternative, forced choice task. **C**, Dataset 2: Electrocortigraphy (ECoG) from nine human subjects were analyzed as they performed a finger flexion task [13]. **D**, Electroencephalography (EEG) systems with 129 - channels were used to record brain-wide activity from 9 human subjects performing unconditioned stimulus task.

To evaluate this atomic decomposition as a denoising technique for field potentials, we utilize our previously proposed transient model for phasic event extraction [10]. The advantage of the generative model lies in its ability to characterize with high-resolution, the timings and amplitudes of burst events in bandpassed traces of single-channel recordings corresponding to the clinically established frequency ranges (delta to high gamma). The statistical nature of the technique and the employed optimization ensure that the detections strictly adhere to the neurophysiological properties of oscillatory bursts. Using the M(PP)-spectrogram defined in [11] to capture fine time variations in the bursting power, we demonstrate that an MPP representation is effective in alleviating spurious linear dependencies between multi-channel recordings, regardless of the recording medium.

Considering that a sparse representation of channel activity will inherently reduce correlations between the channels, we justify our event-based representation by demonstrating its ability to preserve the properties of the underlying neural dynamics - both in function and location within the brain network. Firstly, we validate the retention of behavioral information in the MPP signal by examining beta and gamma activity in the motor cortex during movement initiation and execution. By combining information about beta and gamma bursts in a non-linear decoding model, we achieve successive, asynchronous prediction of the time-points of finger movement and the corresponding finger label. Secondly, we illustrate that an MPP representation maintains effective connectivity information between areas by recovering top-down functional coupling between the motor and prefrontal cortices from traces of high-gamma burst rates.

Overall, our studies confirm that a marked point process representation of single channel activity using oscillatory bursts enables a minimally correlated, high resolution, descriptive representation of electrophysiological signals, surpassing the capabilities of conventional time-frequency methods. Additionally, the neurophysiological basis of this representation allows for seamless translation between the multi-scale observations. Finally, the model itself is a computationally inexpensive tool to further understand the behavioral and functional implications of region-specific field-potentials.

## Materials and methods

### Datasets

*Dataset-1* : Local field potentials were recorded from the dorsal prelimbic cortex (dPrL) of three rats performing a two-alternative, instructed delay forced choice task [12]. A 32-channel microwire array was implanted in the right hemisphere, with 16 electrodes positioned in the medial prefrontal cortex (mPFC) and 16 electrodes in the secondary motor cortex (M2).The recordings were downsampled to 500 Hz for analysis.

The experiment was set up in an acoustically isolated operant conditioning chamber with three nose poke holes: a central fixation hole and flanking target holes(Fig.1**B**). Trials were initiated by the subject by positioning its snout in the fixation hole. After a 1s fixation period, the instruction cue was sounded - a variable pitch single tone delivered at 60 dB whose pitch determined the target hole. A low-pitch tone indicated the right target hole, while a high-pitch tone cued the left target hole. The cue initiated a pseudo-random delay period (uniform distribution, **U**(1.5, 2) seconds). A ‘Go’ cue, consisting of a white noise auditory stimulus delivered at 60 dB, signaled the end of the delay period, prompting the subject to place its nose in the correct target hole.

Successful visits were rewarded with a 45mg food pellet (Bio-Serv, NJ), while incorrect visits received no reward. All animal procedures were reviewed and approved by the Institutional Animal Care and Use Committee (IACUC).

*Dataset-2* We analyzed ECoG recordings from a publicly available dataset [13], comprising nine subjects implanted with platinum arrays (Ad-Tech Medical Instrument Corporation, Wisconsin, USA) with a variable number of channels per subject. Recordings were made from various brain regions, including the dorsal primary motor cortex, dorsal primary sensory cortex, and ventral sensorimotor cortex, while the subjects performed a finger movement task (Table S1). Task instructions were conveyed through displayed words, prompting self-paced movements in a 2-second interval followed by a 2-second rest period where subjects fixated on a blank screen. Each finger was moved between two and five times during the 2-second movement periods, recorded using a five-degree-of-freedom data glove sensor (5 dt, Irvine, CA). Approximately 30 movement cues were presented for each finger, with trials interleaved randomly(Fig 1**C**).

The ECoG data were recorded using the *Synamps 2* biosignal amplifier (Compumedics Neuroscan, North Carolina, USA) at a sampling rate of 1kHz, and later were bandpass filtered between 0.3 to 200 Hz. Finger movements were not time-aligned to a particular start time; however, subjects could initiate movement at any point during the 2-second movement period.

Patients participated voluntarily, providing informed written consent under approved Institutional Review Board protocols at the University of Washington (#12193). All patient data was anonymized according to IRB protocol, in accordance with HIPAA mandate. This dataset was originally introduced in the publication, “Human Motor Cortical Activity Is Selectively Phase-Entrained on Underlying Rhythms,” published in PLoS Computational Biology in 2012 [13].

*Dataset-3:* A total of 129 EEG channels were recorded from 20 subjects engaged in a task where an unconditioned stimulus (UCS) was randomly interleaved with a conditioned stimulus (CS) in each trial (Fig. 1**D**). Trials began with subjects fixating on a centrally presented cross on the screen for approximately 3.6 seconds. The conditioned stimuli, a Gabor patch with a 1.5-degree left tilt and a Michelson contrast of 0.63, were then presented for 2.5 seconds. The unconditioned stimulus, a 96 dB sound, was randomly paired with the CS in about 50% of trials and occurred approximately 1.5 seconds after trial initiation, lasting for 1 second. The epochs analyzed for this study were 5.1 seconds in duration, including a 3.6-second display of a fixation cross before trial initiation and 1.5 seconds after CS onset. The data were sampled at 500 Hz and subsequently averaged to a single reference. The study received approval from the local Institutional Review Board.

### Transient model for phasic events

Oscillatory burst events are a particularly descriptive feature of field potentials. Appearing as sparse, transient and ordered patterns amidst the background activity, they stand out for their periodic time structure and significantly high power. Given their properties, these transient events have been suggested to serve as a communication channel between adjacent neural populations [8]. The background activity, on the other hand, follows the 1/f power law and exhibits an approximate Gaussian distribution in its amplitudes [8]. Expanding on this two-state hypothesis, the transient model for phasic events incorporates frequency-specialized information derived from established cortical rhythms. In this model, single-channel, bandpassed traces of neural recordings, 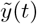, are defined as a linear combination of background activity, *n*_0_(*t*) and oscillatory bursts, 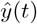 (1). Next, each burst event is modeled as the impulse response of a linear filter centered at a particular frequency (*ω*) within the clinically established frequency range (2). These set of filters or dictionary atoms, 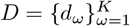, represent ‘typical’ oscillatory-responses in the bandpassed traces. Finally, 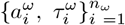 are the projection coefficients and the timestamps of the burst events best described by the filter *d*_*ω*_.

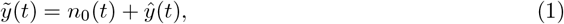

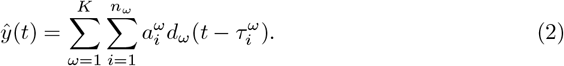

The descriptors of each oscillatory burst, i.e., its time of occurrence (*τ*_*i*_), maximum amplitude (*α*_*i*_), specific narrow-band frequency (*ω*) and duration (*δ*_*i*_) constitute a Marked Point Process (MPP); wherein lies the model’s advantage to characterize properties of oscillatory bursts with a time resolution given by the sampling period, defined for each of the clinical frequency bands.

To learn such a representation, our description of the field potentials uses dictionary learning at multiple frequencies, instead of the more conventional spectral techniques. The model is realized in two phases: 1) a denoising phase that identifies segments with burst events, and 2) a learning phase that constitutes the blind estimation of the filter bank, *D*, from the training data. Since the dictionary atoms are obtained directly from the filtered, noisy field potentials, the ‘denoising’ phase uses a correntropy based clustering technique to discriminate between the background activity and burst patterns [14]. In the histogram of amplitudes within the selected clinical frequency range, the deviation from the Gaussian distribution at the high amplitude tail, indicates the mixing between background activity and the oscillatory bursts. To determine this boundary, the method iteratively selects and adjusts a threshold *κ* using the full distribution information of the bandpassed trace to best separate background activity from oscillatory bursts. Only the time segments of length M that have power above *κ* are included as putative bursts in the learning phase.

Next, the learning phase performs alternating optimizations between sparse coding and (correntropy - based) dictionary-learning. This process iteratively selects segments for the dictionary, incorporating constraints on the minimum dictionary length and minimizing the reconstruction cost of the bandpassed traces (2). Model training is terminated when any one of the convergence criteria are achieved: reaching a specific number of iterations or a saturation on the Frobenius difference norm between dictionaries from successive iterations. Once the dictionary is learned, the construction of the MPP feature space involves a simple convolution between the learnt filter bank and the test data. When the convolution output surpasses the threshold *κ* for any one of the learnt filters, a burst event is identified at that particular time stamp.

### Parameter selection

The model relies on two hyperparameters: the maximum duration of the oscillatory burst event, *M* and the number of filters, *K*, in the filter bank. The hyperparameter *M* dictates the upper limit on the transiency of the burst events within the specified frequency range. In this study, the value of *M* for beta and (low/high) gamma burst events are set as 500ms and 100ms, respectively. These values primarily follow from the neurophysiological principles of the cortical rhythm that we further verify via a thorough visual inspection of the bandpassed traces in the time-domain. Next, the maximum cardinality of the learnt dictionary is decided by the hyperparameter, *K*. For both frequency ranges, this cardinality is upper bounded to *K* = 30 filters, as the number of detected burst events seemed to saturate close to this limit (tested on training dataset for *K* = *K*′ × *M, K*′ = [0.125 − 1], step-size = 0.125). Lastly, both denoising and dictionary-learning phases incorporate the correntropy measure [14], which requires selecting the bandwidth of the Gaussian function using Silverman’s rule [15].

### M-spectrogram, M-rate

An MPP representation enables the construction of a rich feature space via appropriate combinations of the marked features by evoking measures and methods from the well-established field of point process systems. The M-spectrogram and -rate are two such constructs [11]. While the M-rate (*λ*) is the traditional intensity function of the point-process that is estimated via kernel smoothing, the M-spectrogram is an extension to the intensity function in that it is weighted by the transient power within the oscillatory burst at its time of occurrence. Given the timings of each burst event, *τ*_*k*_ : *k* = 1, 2, …*N*, the construction of the M-spectrum is summarized in (3)–(5). First, transient burst power, 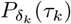, is estimated as the norm-square of the signal amplitude values in the duration of the event, (3). Next, the power-weighted MPP events are aligned to the onset of the stimuli and superimposed to form the marked event density, 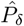, (4). Finally, an estimate of the local burst power, i.e., the M-spectrogram (*λ*_*α*_), is obtained via Gaussian kernel smoothing (5).

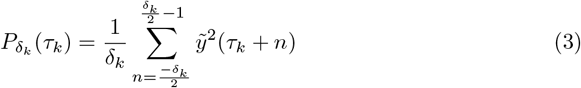

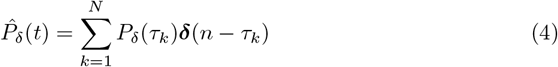

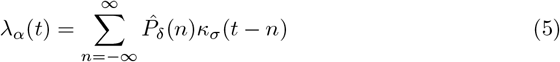

The M-spectrum was specifically designed to address a major shortcoming of the time-frequency (TF) methods - the inherent trade-off between frequency and time resolution in Fourier decompositions. Unlike TF methods, the transient model does not impose any predetermined structures on the signal and instead leverages time information within the signal to construct the dictionary. Therefore, the M-spectrum, solely built on the power in the oscillatory bursts preserves time information in the frequency range of interest while consistently limiting leakage from power due to background noise.

The M-spectrograms and -rates are estimated from a point process representation using Gaussian kernels and it is important that the selection of its bandwidth be neurophysiologically informed. For all our studies, we chose a bandwidth size that approximately covers the burst duration, that translates to 50 and 30 ms for beta and gamma burst, respectively.

### Data Preprocessing

We processed data consistently across all subjects (both human and non-human) and sources (EEG, ECoG, and LFP), unless specified otherwise. To mitigate power-line artifacts within the frequency range of interest, IIR Notch filters were applied, and all data were uniformly downsampled to 500 Hz. Each trial was aligned to the stimulus onset, and the complete trial duration was utilized for analysis. For the online movement classification task (Figure 1**C**) and preceding analyses involving ECoG data, an additional common-average referencing was employed to minimize signal bias towards any specific scalp location. Before applying the transient model, channel activity was filtered using a zero-phase FIR bandpass filter within the frequency range of the cortical rhythm. To explore the dynamics of oscillatory bursts in the beta and gamma bands, four filters were designed using Hamming windows with a quality factor, Q*∼*1, and specific center frequencies (Table 1). Throughout our analysis, comparisons with Short-Time Fourier Transform (STFT) were incorporated. In all analyses, STFTs were computed using a window size of 0.5 seconds and a 50% overlap between windows.

**Table 1.** Parameters of FIR bandpass filters in clinical frequency ranges.

| Band | Center<br>frequency<br>(Hz) | Order |
| --- | --- | --- |
| Beta (Rat<br>LFPs) | 20 | 43 |
| Beta (Human<br>ECoG) | 30 | 40 |
| Low-Gamma | 60 | 33 |
| High-Gamma | 115 | 25 |

### Asynchronous online classifier for motor events

For decoding motor movements at the level of individual fingers (Fig 1**C**), we exploit the temporal resolution of the multi-channel M-spectrum to improve classifier accuracy (Fig 2**A**). We construct an asynchronous online classifier comprising of two stages: (1) a movement detection stage to predict the precise time-points of the motor movement (Fig 2**A**(middle)), and (2) a digit classification stage built solely on the multi-channel spectral information during movement (Fig 2**A**(right)). By using this approach, we do not require information about the time of the stimulus, unlike many competitive algorithms. Finally, the results from the hierarchical classification are combined to interpret the final finger-level movements. Below, we provide specific details about the classifiers constructed for each stage.

**Fig 2.**
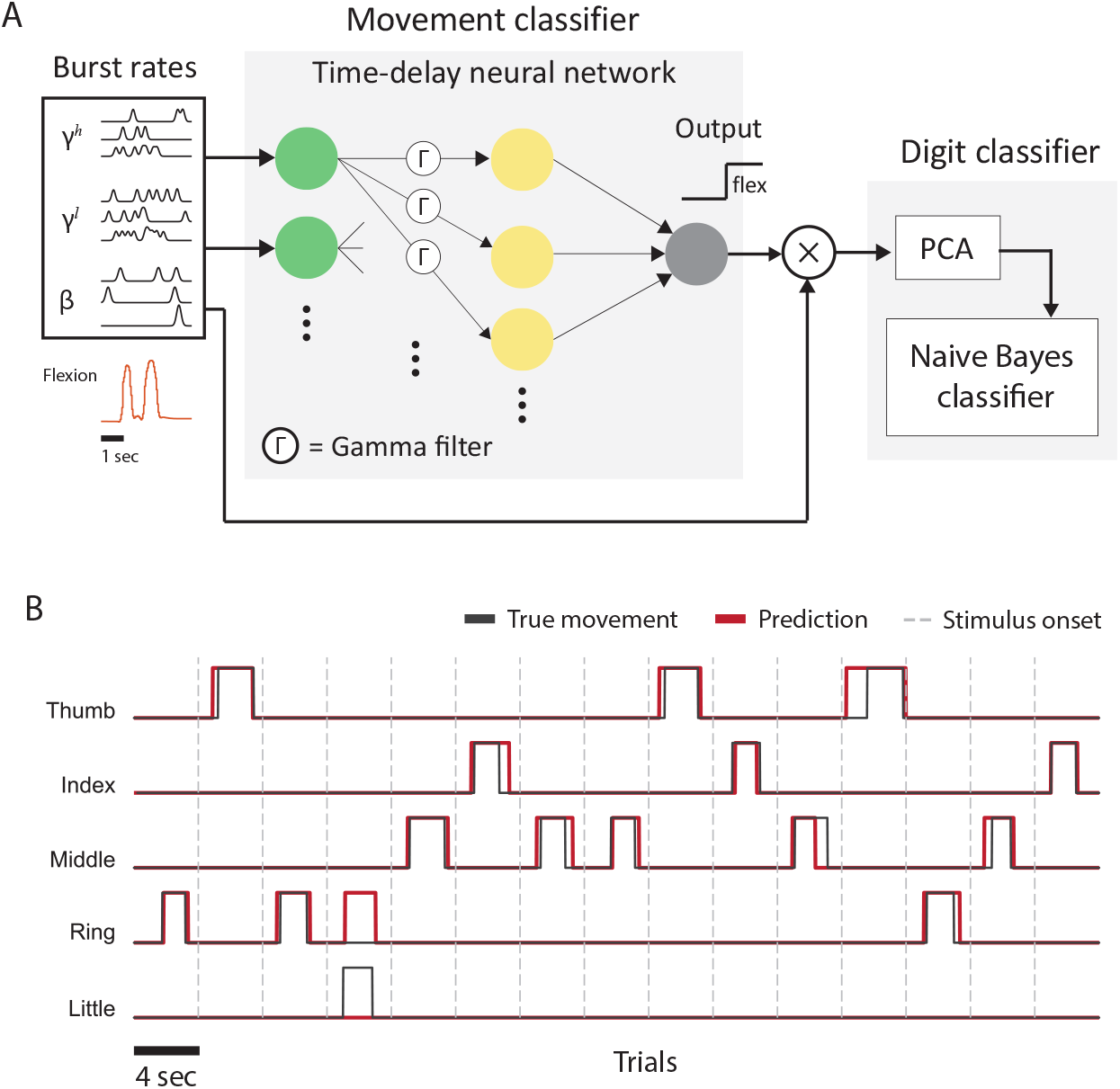
Asynchronous online classifier for motor movements. **A**, The schema of the online asynchronous classifier consists of separate modules for movement classification (middle panel) and digit classification (right panel). The variables *β, γ*^*l*^, and *γ*^*h*^ represent the M-spectra in the beta, low-gamma, and high-gamma bands, compiled across channels, which serve as inputs to the classifier. The movement classifier is a single-layer perceptron equipped with a gamma memory filter (Γ) that predicts the time points of movements. The mean burst rates during these movements are then evaluated based on the predictions from the movement classifier and passed to the digit classifier. The digit classifier projects the burst rates across all channels onto a lower-dimensional space using principal component analysis (PCA) and then applies a Naive Bayes classifier to predict the specific finger associated with the movement. **B**, A 60-second snippet demonstrating classifier performance in 15 trials (Subject 1).

### Movement detector

To extract temporal movement information from the multi-channel M-spectrums, we aimed to incorporate a time-series classifier. While advanced options like long-short term memory classifiers and recurrent neural networks are common choices, for the purpose of demonstrating the effectiveness of M-spectrogram for behavior prediction, we opted for a simpler classifier – time delay neural network (TDNN) built using a gamma memory and a single layer perceptron. This classifier can be implemented in ambulatory subjects due to its simplicity and compatibility with ultra-low-power microprocessors. The gamma memory unit serves as an improvement over the tapped-delay line framework frequently used in neural networks [16]. The unit is a single-input multi-output filter that is a cascade of *L* leaky integrators with the same time-constant *μ*. The value of *μ* dictates the system memory depth and the frequency response of the filter. Choosing 0 *< μ <* 1 generates a low-pass response, while 1 *< μ <* 2 results in a high-pass filter response at each tap. The gamma memory unit in the classifier network is a 15^*th*^ order filter with a time dependency defined by *μ* = 0.2.

Outputs from the gamma memory unit are fed into the input layer of the MLP. The MLP comprises of a single hidden layer of 30 processing elements that implement a rectified-linear activation function (ReLu). Learning on the model includes maximizing a correntropy cost function [17] using backpropagation with the adaptive moment-estimation method, adam [18]. A movement classification threshold was applied to the model’s predictions to classify each time point into either a ‘moving’ or a ‘resting’ state. After this, to suppress the very short on-off detections, a smoothing window of size, *W* (*∼*0.8 − 1.2*s*), was employed. Finally, the smoothed movement output along with the input M-spectrums were passed as inputs to the finger-label classifier.

### Finger-label classifier

Identification of the movement time-points allows access to the locally relevant transients in the cortical rhythms via the M-spectrum. Features for the finger-label classification were then constructed as the mean of the M-spectrum in the time-interval [−0.25 *− t*_*∞*_]s, where, ‘0’ and ‘*t*_*∞*_’ mark the predicted time-points of movement onset and offset, respectively. Ideally, a high-resolution spatial distribution of the channel activity would reveal clear discriminative patterns between finger labels [19]. However, in the absence of such resolution, we resort to principal component analysis (PCA) to differentiate the most variable dimensions among the channel features. We do so while preserving 98% of the explained variance among the channel features. The reduced outputs from PCA are then used for label classification via a cross-validated, regularized linear discriminant analysis (LDA). Regularization helps improve generalizability by weighting the non-diagonal matrix elements of the within-class scatter matrix, *S*_*w*_, that may contribute to an overfit classifier. Specifically, *S*_*w*_ is replaced with a regularized estimator, *S*_*wr*_ = (1 *− p*)*S*_*w*_ + *p***I**, where **I** is the identity matrix, and *p* is the cross-validated shrinkage parameter.

### Performance Quantification

For each subject, we performed a 5-fold cross-validation over all trials and quantify the performance of both classifiers as well as of the combined system by means of accuracy. For both the movement (Table 3, column 1) and the final classifications (Table 3, column 3, Fig 5**B**), accuracy is calculated as the proportion of correctly classified labels for each time-point averaged over all trials (i.e., 0*/*1 for movement classification and 0*/*finger-digit for final classification). The accuracy for finger-label classification (Table 3, column 2) is calculated as the ratio between correctly classified trials and total number of trials. Further, receiver operating characteristics (ROCs) were analyzed for each window, *W*, and movement classification threshold over the movement classification, where sensitivity and specificity are evaluated as per (6) and (7), respectively (Fig 5). The window and threshold best suitable for classification were chosen by identifying the best classifier accuracy on the complete task (marked by the golden marker in Fig 5**A**). Finally, for a complete summary, we also analyzed the area under the curve (AUC) of each ROC (Fig 5**A**, inset).

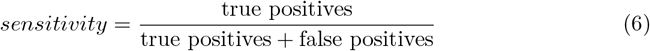

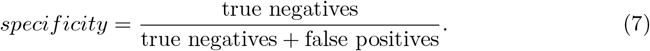

### Effective connectivity across brain areas

To check whether the MPP representation can accurately recover neuronal interaction patterns in recorded LFP activity (Fig 1**B**), we used directed information to quantify effective connectivity. Directed information (DI) captures the feedforward flow of information between two time series [20]. For two variables, *X* and *Y*, directed information is formally defined using entropy as shown in (8). Here, *X*^*n*^ denotes the successive samples of the process, *X*, between time-points 1 and n, i.e., *X*^*n*^ = [*X*_1_, *X*_2_, …, *X*_*n*_]. The term *H*(*Y* ^*n*^|*X*^*i*^) is called ‘causal conditional entropy,’ distinguishing itself from conventional conditional entropy by restricting the samples of *X* to *i* rather than extending over the entire dataset *n*. This restriction makes DI a causal measure of information flow. Lastly, in the absence of a causal influence of *X* on *Y*, the causal conditional entropy reduces to an entropy formulation and therefore, *I*(*X*^*n*^ *→ Y* ^*n*^) = 0.

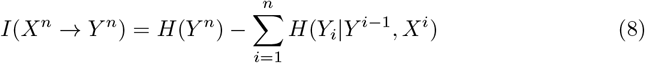

The computation of directed information typically requires knowledge of the joint probability distribution function, which is often unknown in neuroscientific scenarios. In our study on effective connectivity, we adopt an approach circumventing direct estimation of the probability distribution. Instead, we leverage a Hilbert space approach utilizing the matrix-based Renyi’s *α*-order entropy, as proposed by [21]. This measure defines entropy over the eigenspectrum of the normalized Gram matrix, a Hermitian matrix derived from the projected data in a reproducing kernel Hilbert space (RKHS). For a random variable *X* = *x*_1_, *x*_2_, …, *x*_*n*_ and a Gram matrix *K* evaluated on *X* using the real-valued positive definite kernel *κ* : 𝒳 × 𝒳 ⟼ ℝ, the matrix-based Renyi’s *α*-order entropy is given as shown in (10), where 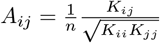 and *λ*_*i*_(*A*) denotes the *i*^*th*^ eigenvalue of *A*.

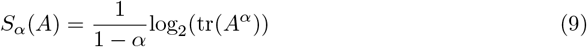

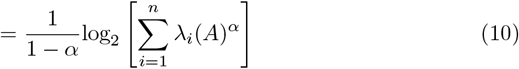

The sample data mapped into the Reproducing Kernel Hilbert Space (RKHS) via the induced kernel allows computation of higher-order statistics. Variations in these moments are then quantified through the eigenspectrum of the estimated Gram matrix. Using the above estimator of entropy, we obtain the following expression for directed information,

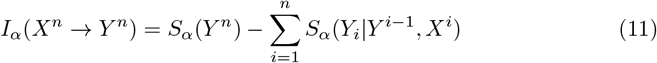

While our approach efficiently formulates directed information, defining a positive definite kernel for point processes without a natural algebraic structure remains a challenge. Focusing on kernel-based methods for structuring neural spike train data [25, 26], we adopt concepts from [27]. Firstly, the point processes are brought into the continuous *L*_2_ space as intensity functions via convolutional operators. Next, leveraging an RKHS-inducing kernel, *κ* : 𝒳 x 𝒳 ↦ ℝ, we apply the kernel trick. This involves subjecting the intensity functions to pairwise kernel evaluations. The resulting Gram matrix, *A*, is a symmetric positive semidefinite matrix representing pairwise inner products in the Hilbert space. In this way, Kernels introduce structure into the point process space, with the Gram matrix capturing higher-order information. Notably, directed information estimation tends to yield a biased estimation between independent time-series. Therefore, to identify true connections, we implement significance testing using a null distribution based on a shuffled time series.

Our formulation of directed information depends on three hyperparameters: 1) the kernel that determines the conditional intensity function, 2) RKHS-inducing kernel, and 3) the order *α*. To evaluate the conditional intensity, we use a 60 ms rectangular window function. For the RKHS-inducing kernel, we have identified the non-linear Schoenberg kernels as the most suitable choice for computing the causal measure [28]. The kernel formulation for two intensity functions, *x* and *y*, is shown in (12), where the intensity functions are formulated as in (13) such that 𝕀 is the indicator function, *n*_*w*_ is the window size and 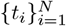 are the event timings. The kernel size, *σ*, is determined by Silverman’s rule for density estimation [15].

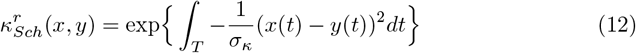

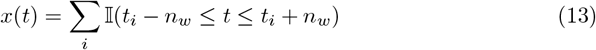

Lastly, the choice of *α* is associated with the task under study. For emphasis on rare events (i.e., in the tails of the distribution), *α* must be chosen close to 1, while higher values of *α* (*>* 2) characterize modal behavior [21]. Since in our analysis of effective connectivity, the focus is on the sparse transient synchronizations between channel activity, the order *α* is set to 1.01.

## Results

To assess the effectiveness of an MPP representation in attenuating volume conduction effects, we conducted a comprehensive analysis across three sensing modalities: Electroencephalography (EEG), Electrocorticography (ECoG), and local field potentials (LFP). This approach allowed us to explore and compare neural dynamics across diverse behaviors, species, and time-frequency models.

### Improving effects of volume conduction

Oscillatory bursts represent the source-specific patterns of channel activity, reflecting short-lived synchronizations between neural networks across various spatial scales. Given their prominence at each scale, specific statistical properties, and frequency specialization, we hypothesize that an oscillatory bursts-based representation of electrophysiological signals will aid in limiting the spatial and temporal extent of volume-conducted sources. To study the influence of volume conduction on these burst events, we examine inter-channel correlations in signals recorded across EEG, ECoG, and LFP modalities. The M-spectrogram, focusing on the oscillatory bursts’ signal power, serves as a key metric to test our hypothesis. Throughout our analyses, we compare the performance of the M-spectrogram against standard measures such as STFTs and instantaneous power in the raw signal.

First, we evaluated the percentage decrease in zero-lag inter-channel correlations obtained using M-spectrum (and STFT) relative to the instantaneous power in the raw signal. To explore the spatial impact of this reduction, we analyzed power spectra in both low and high-frequency ranges—specifically, in the beta band (ECoG, EEG: 20-40 Hz, LFP: 10-30 Hz) Fig S1) and the gamma bands (EEG: 40-80 Hz, ECoG, LFP: 80 - 150 Hz; Fig 3**A-C**). For each trial and channel pair, we evaluated the Pearson correlation coefficient between the estimated signal power over the complete duration of the trial (Fig 1**B-D**). On the resulting trial-averaged correlation matrix of size, number of channels × number of channel, applying a column mean presents the extent of linear dependence between channels over the brain surface. To account for the spatial resolution of EEG and ECoG signals, we investigated both local (Figs 3**A-C**, S1(bottom)) and global (Figs S1, S2(top)) correlations in these modalities. Local correlation patterns were examined using six probes in the nearest vicinity of each channel, while global correlations were assessed with the farthest twenty channels for EEG and ten channels for ECoG. For LFPs, average correlations were calculated between all channel pairs, considering their local spatial extent.

**Fig 3.**
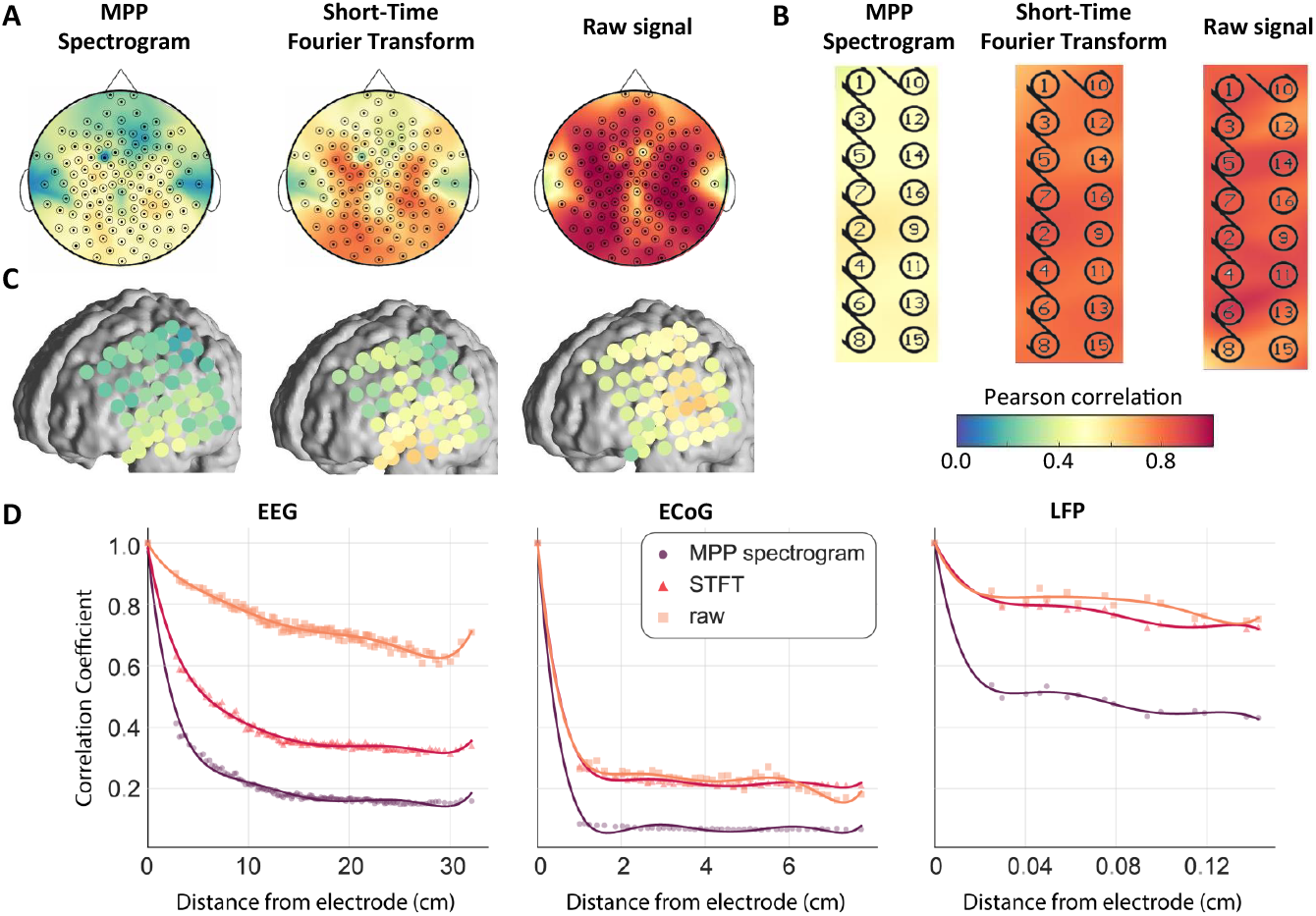
Comparison of local cross-channel correlations between high-gamma oscillations across different sensing modalities and models. **A**, Average Pearson correlation coefficient of each EEG channel with its 6 nearest neighboring channels, evaluated on (left) a marked point process (MPP) representation of bursts, (middle) short-time Fourier transform (STFT), and (right) raw EEG for an example subject. **B, C** The same analysis as in **A**, applied to (**B**) LFPs recorded in the prefrontal cortex of a rat and (**C**) ECoG data from the left fronto-parietal area in a human, respectively. Data shown is from Subject 1 in each dataset.**D**, Correlation as a function of distance from each channel across sensing modalities, calculated for the same subject as in **A–C**. Correlations are presented as the average Pearson correlation coefficient between pairs of channels, with distances measured as the Euclidean distance between channels. Markers represent the average Pearson correlation between channel pairs and solid lines establish the best-fit lines (obtained using polynomial curve fitting, n = 5), capturing the correlation trends with distance.

Our initial findings revealed significant reductions in inter-channel correlations within M-spectrograms across all three observation scales, averaging approximately 45% (Table 2, Figures 3**A-C**, S1, S2). Importantly, we observed higher reductions in global correlations compared to local correlations (p *<* 0.001, ANOVA). LFPs exhibited the smallest decrease in correlations, whereas ECoG and EEG channels demonstrated a comparable reduction of about 50% in inter-channel correlations (p = 0.54, ANOVA). For EEG and ECoG, M-spectrums in the gamma range exhibited lower correlations across channels compared to those in the beta range (p *<* 0.001, ANOVA). In contrast, STFTs showed a significantly smaller decrease (p *<* 0.001, ANOVA) in the linear dependencies between channel activities. Despite M-spectrograms being less correlated across channels, their spatial correlation patterns aligned with those of the STFT and instantaneous power.

**Table 2.**
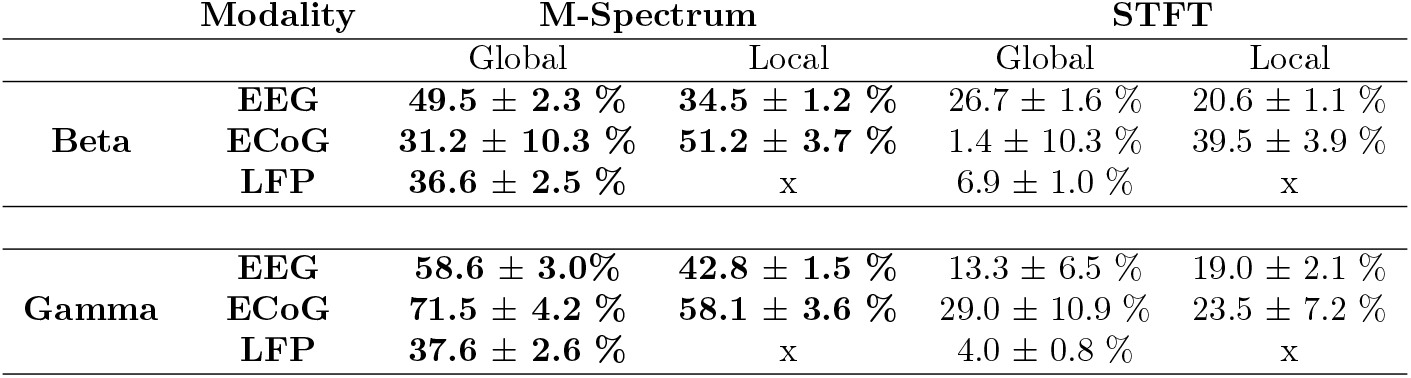
Average percentage decrease in inter-channel correlations relative to raw signal power.

**Table 3.** Subject-wise performance of the online finger-level movement classifier at each classification stage.

| ID | Movement<br>Classifica-<br>tion<br>(2-class) | Finger<br>Classifica-<br>tion<br>(5-class) | Final<br>Accuracy<br>(6-class) |
| --- | --- | --- | --- |
| 1 | 86.4 ± 1.4 | 75.1 ± 2.2 | 76.3 ± 1.5 |
| 2 | 86.4 ± 2.3 | 78.0 ± 2.7 | 73.3 ± 2.0 |
| 3 | 79.1 ± 1.5 | 79.2 ± 1.7 | 67.0 ± 1.1 |
| 7 | 81.4 ± 1.7 | 78.0 ± 2.7 | 71.1 ± 1.4 |
| 8 | 85.5 ± 0.5 | 69.2 ± 2.1 | 72.4 ± 1.8 |
| 9 | 79.9 ± 1.6 | 80.8 ± 2.3 | 70.0 ± 2.2 |
| Avg. | 82.6 ± 1.6 | 76.7 ± 2.3 | 71.7 ± 1.7 |

To understand the spatial extent of volume conduction effects in each model, we examined the correlations between channel power across varying distances from the electrode for signals in different frequency bands (Fig 3**D**). For each trial, we computed correlations between all channel pairs and averaged them across all trials. To estimate the distance between channels, we computed the Euclidean distance between their coordinates in the 2-D space and scaled each value by the minimum distance (in cm) between two electrodes. After aligning all channels based on their distance, we averaged the correlations, deriving a representative mean correlation over distance. Across the different models, M-spectrums not only noted the lowest correlation but also demonstrated the steepest declines. The reduced effect of volume conduction on M-spectrums was particularly notable in EEG recordings. These findings confirm a substantial reduction in volume conduction effects on the M-spectrogram, surpassing those observed using STFTs.

### Phasic events as descriptors of behavior

Given the significant reductions in both local and global inter-channel correlations, our subsequent goal was to determine whether the reduction is merely a by-product of a sparse representation or a deliberate outcome of our approach that builds on the premise that transient population-level synchronizations serve as carriers of information in time. If the latter hypothesis holds, the features of the marked representation must encode region-specific functional information, such as correlates of pertinent behavior or location within the network hierarchy.

Desynchronization of beta rhythms during motor initiation and production is a commonly observed phenomenon in the somatomotor regions of the cerebral cortex [29, 30]. Additionally, movement in these tasks is associated with an increase in gamma power, exhibiting a more focal cortical distribution corresponding to the regions of the involved populations. Given these findings, we asked if the M-spectrum can capture such cortical phenomena. For this, we studied the MPP feature space of ECoG signals recorded from 9 subjects while they performed a finger flexion task (Fig 1). Throughout the task, field potentials were recorded over various areas, including the primary motor and sensory cortices.

To validate the M-spectrum’s ability to identify motor areas based on their functional role in the movement task, we conducted a channel-specific regression analysis. The goal was to predict finger labels using the mean beta and gamma M-spectrum (both low gamma between 40 - 80 Hz and high gamma between 80 - 150 Hz) in each trial. Across all subjects, the analysis consistently revealed the highest decoding accuracies in channels located in the primary motor cortex (Fig 4**A**), confirming the M-spectrum’s capability to preserve the region-specific functional significance of the brain area (Table S2). Subsequent analyses focused exclusively on channels in and around the motor cortex, following the demarcations in the author’s notes [13] (electrode regions marked between [0 − 3]). In this section, any reference to ‘channels’ implicitly denotes the selected subset within the motor areas.

**Fig 4.**
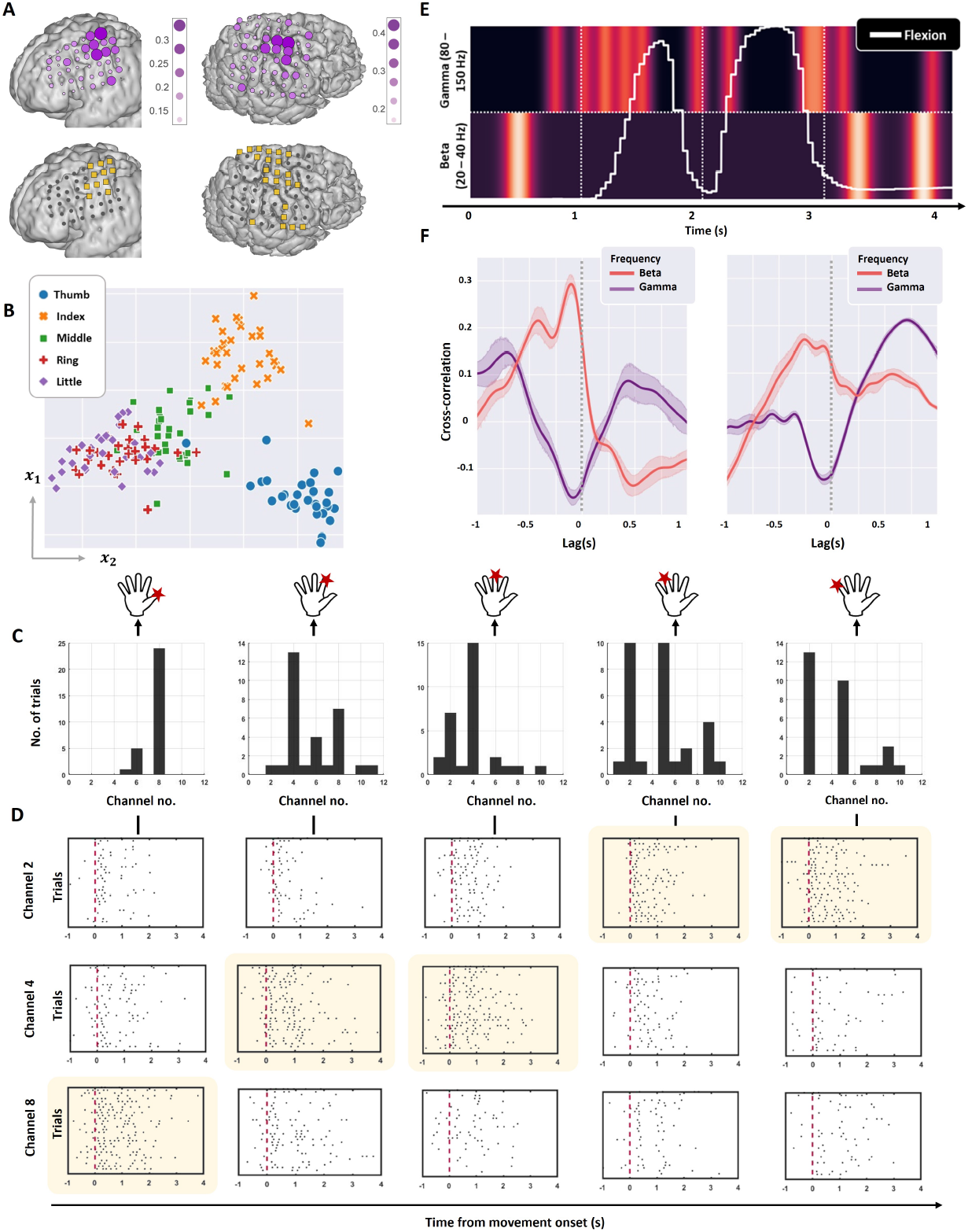
Motor correlates within oscillatory burst rates. **A**, Top: Cross-validated decoding accuracy for predicting finger labels, obtained using logistic regression on the mean gamma (low and high) and beta M-spectra for each ECoG channel in two exemplary subjects (left: Subject 1, right: Subject 2). Darker colors and larger circle sizes indicate higher decoding accuracy. Bottom: For the same subjects, highlighted channels correspond to those located in motor areas. **B** Linear projection of the mean gamma (low and high) band M-spectra during movement into a 2-dimensional space using PCA for an example subject (Subject 3). **C**, Histogram of channels with the highest high-gamma activity per trial, categorized by the corresponding finger moved. **D**, Raster plots of high-gamma bursts from the three ECoG channels showing the highest gamma activity across different fingers in Subject 1. Each plot is aligned to movement onset (red line), with yellow shading indicating the channel with the highest gamma activity for each specific finger. These plots illustrate finger-specific specialization of gamma bursting across channels.**E**, Single-trial M-spectra in the high-gamma (top) and beta (bottom) bands superimposed with finger flexion (white trace) for Subject 1, Channel 3, Trial 45, and the middle finger.**F**, Average cross-correlation of finger flexion with beta (red) and gamma M-spectra (purple), showing movement-specific entrainment between these frequencies for Subjects 7 (left) and 9 (right).

**Fig 5.**
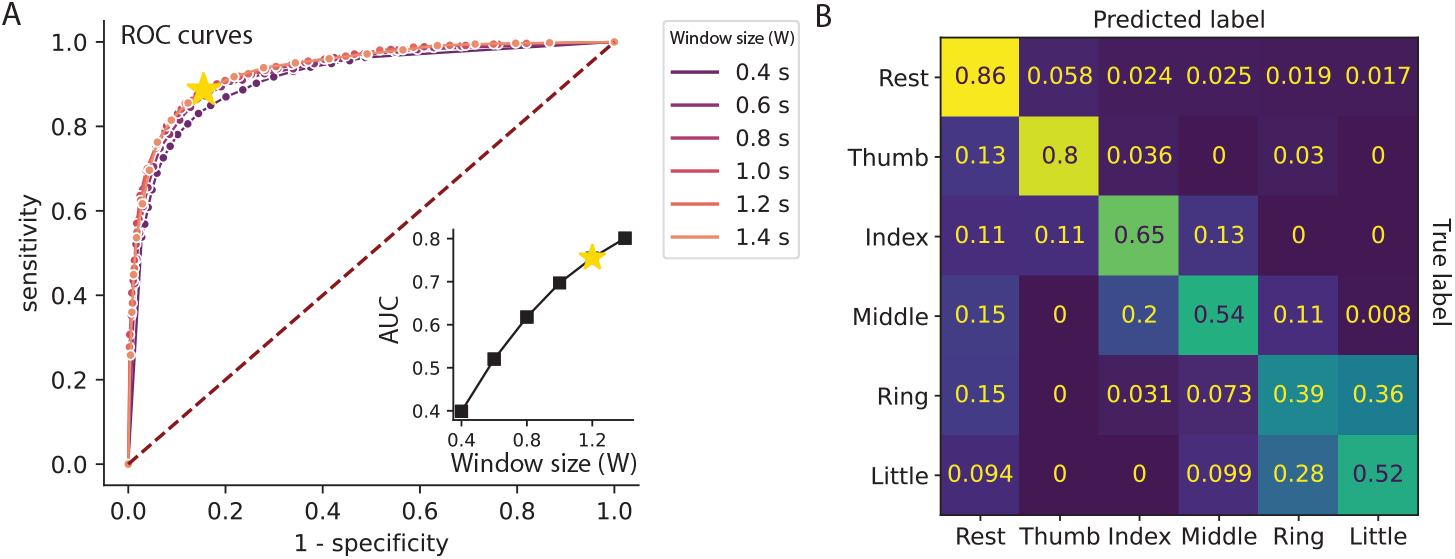
Performance of the online, asynchronous finger-level movement classifier. **A**, Receiver operating characteristics (ROC) curves for multiple windows obtained by varying the threshold for movement classification. Inset: For each ROC curve, the corresponding area under the curve (AUC) is presented. **B**, Example confusion matrix for the final 6-class classification (window size, *w* = 1.2s, movement classification threshold = 0.6). All results are obtained from Subject 1.

Next, we were interested in analyzing the specialization of neural populations in controlling finger-specific movements as captured by the M-spectrum. For each trial, we identified the channel with the highest mean high-gamma M-spectrum in the interval [− 0.25*− t*_*∞*_]s, where ‘0’ and *t*_*∞*_ denote the movement onset and offset times, respectively. Grouping trials by the finger label, we constructed a histogram illustrating the frequency of channel selection for each finger (Fig 4**D**). The histogram revealed finger-specific channel activations across all channels. Next, we visually examined raster plots corresponding to high-gamma activity in three channels, each corresponding to a channel selected most frequently per finger (Fig 4**D**). Aligned to the time of movement onset, the raster plots revealed sustained high-gamma activity in the most-picked channel during finger movement, while high-gamma activity in other channels was sparser and of lesser power. To further substantiate our observations, a 2-component principal component analysis (PCA) on the mean gamma M-spectrum during movement demonstrated separability across distinct finger labels (Fig 4**B**). Across analyses, the overlap between the little and ring fingers is worth noting where the movement of one usually causes movements in the other [19]. Hence, M-spectrums effectively preserved the neural dynamics reflecting behavior, even at the sub-level of individual fingers.

To assess the entrainment between beta and gamma activity as captured by the M-spectrum [29, 30], we visually aligned finger flexion with M-spectrums in both frequency bands (Beta: 20-40 Hz, high gamma: 80-150 Hz, Fig 4**E**). Such visualization revealed increased high-gamma activity that accompanied a lack of beta bursting during movement. To systematically quantify this relationship across all trials, we conducted a cross-correlation analysis. First, we grouped M-spectrums in the beta and high-gamma bands, along with the respective finger-flexion movements, based on finger labels. Following this, we computed the average cross-correlation of the finger flexion separately with the high-gamma and beta M-spectrums in a finger-specific manner over lags in the interval [− 1, 1]*s* (Fig.4**F**). While the anti-correlation between the beta and high-gamma spectrums is evident, it is also worth noting that the movement initiation is marked by an increase in gamma activity *∼*0.5s post movement, preceded by a decrease in the beta activity prior to movement onset.

To further validate the behavioral information retained in the MPP representation, we sought to construct an online, asynchronous finger-level movement classifier. Our observations about the neural dynamics captured by the model informed its construction. First, we leveraged the observed entrainment between beta and gamma rhythms during motion (Fig 4**F**) to identify the precise timing of movement initiation as well as its duration. Subsequently, recognizing that regions in the motor cortex corresponding to the moved finger exhibit higher sustained gamma activity than others during movement (Fig 4**C, D**), we chose the spatial distribution of gamma power post-movement initiation as a suitable feature set for finger-label classification. However, our analysis of the specificity of channels to the finger-label revealed many overlapping electrodes. We attribute this to the poor spatial resolution of the standard ECoG electrodes and recognize that spatial distribution of gamma activity as a feature for finger-label classification would be more discriminative in high-density ECoG [19]. Despite this limitation, we attempted to compensate for the lack of such resolution via feature reduction to reveal the most discriminative and variable dimensions of the feature space (Fig 4**B**).

Consequently, we constructed a two-stage online classifier for successive finger movement and label classification (see Methods, Fig 2**A**). M-spectrums in the beta, low- and high-gamma bands estimated over the entire trial duration were given as inputs to the classifier. While the first stage of the classifier predicted the onset and execution of movement, the second stage predicted the finger label based on activity around the predicted movement onset. Movement classification in the first stage implemented a time-delay neural network using a gamma memory unit [16]. This addition introduces short-term memory into the neural architecture without increasing the hyperparameter space. Finger-label classification is then performed on the top principal components of the input features using a Naive Bayes classifier. Across six subjects, the model achieved an overall accuracy of approximately 72%, where the prediction accuracy for movement onset and execution was 82.6 ± 1.6%, and the accuracy for finger-label prediction was 76.7 ± 2.3% (Table 3, Fig 5**B-D**). Results from subjects 4, 5 and 6 were excluded due to insufficient training data and poor coverage of the motor cortex (Fig S3). Despite using fewer features and simpler classifiers, these accuracies are comparable to state-of-the-art online machine learning classifiers [19, 32, 33]. These findings underscore the effectiveness of an MPP representation in preserving behavior-relevant information and providing exceptional time resolution for deciphering millisecond changes in behavior.

### Phasic events as descriptors of effective connectivity

Neuronal oscillations are considered a fundamental mechanism for dynamic coordination in the brain [8]. However, the quantification of these interactions is challenged by interpretational caveats, particularly those related to volume conduction. While we have demonstrated that an MPP representation significantly mitigates the impact of volume conduction, it is not obvious whether the model can accurately recover neuronal interaction patterns in recorded activity. In this section, we address this issue through an empirical analysis focusing on the well-documented top-down connectivity between the motor and prefrontal cortices. Studies on rodents [22], monkeys [23] and humans [24] have implicated a top-down influence of neurons in the medial prefrontal cortex (mPFC) over the motor system. Functional coupling between the areas has been proposed as a mechanism to inhibit inappropriate responses based on the set task rules [31]. Since high gamma activity (80 - 150 Hz) serves as a link between single-unit physiology and mesoscale oscillatory dynamics, we study the causal effects between high gamma MPP features estimated from simultaneous recordings of LFPs in the mPFC and secondary motor (M2) areas of the rat cortex (Fig 1**B**).

As the subjects actively engaged in a two-alternative, forced choice task, delay period activity in the two areas revealed peaks in the high-gamma activity that appeared to occur almost simultaneously (Fig.6**A**). Synchrony between gamma activity in both areas was evident not just during the peak gamma rate but also during the shorter gamma events.

To reveal the dominant direction of influence between the areas during the delay period, we evaluated the directed information bidirectionally, estimating *DI*(*mPFC → M* 2) and *DI*(*M* 2 → *mPFC*) on each pair of channels corresponding to the two areas. In trials where subjects performed the task correctly, we observed that the channel-averaged DI values were higher for the direction from *mPFC → M* 2. However, trial-to-trial variations in the M-rate created large variances in the DI estimates. To address this, we performed per-trial normalization of the DI measures across both directions and all channel pairs. These normalized DI measures revealed a significant increase in the DI estimate for the feedforward direction (*mPFC → M* 2) compared to the opposite direction (p *<* 0.05). These results held true regardless of the subject’s final movement direction (Fig 6**B, C**). Combined with the outcomes of the preceding section, these findings underscore the novelty of utilizing oscillatory bursts as a succinct representation encapsulating both encoded behavioral and causal information.

**Fig 6.**
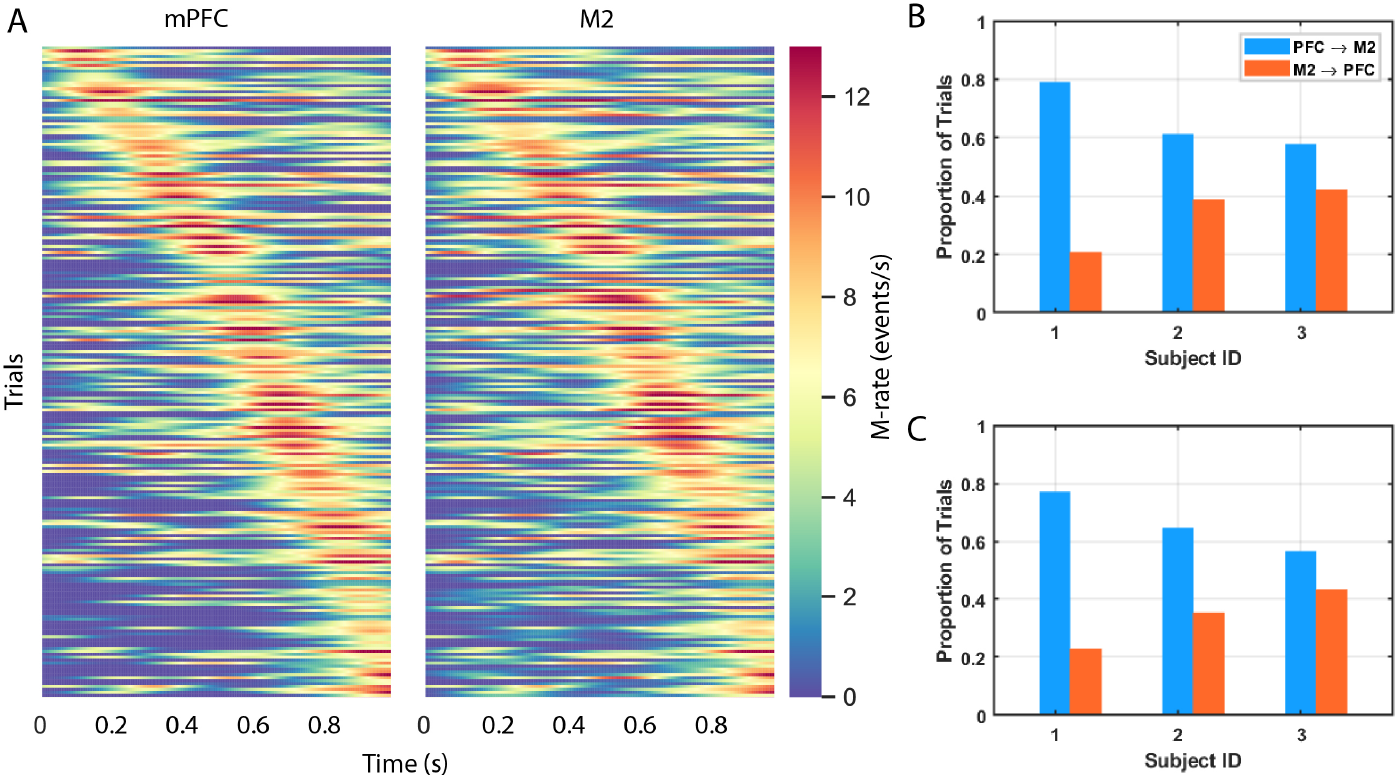
Functional connectivity between secondary motor area and prefrontal cortex in rats. **A** High - gamma M-rate in mPFC (left) and M2 (right) areas, Subject 3, Session 2. Trials are sorted by peak-time in mPFC gamma activity and each row corresponds to the same trial across areas. **B, C** Proportion of trials indicating a causal influence from mPFC → M2 (blue) and M2 → mPFC (orange) i.e., a greater value of DI estimate in the particular direction corresponding to **B** contralateral and **C** ipsilateral movement and instruction. Two sample KS-test revealed significantly higher DI estimate for the direction *PFC → M* 2 in each subject (p *<* 0.05, two sample KS test).

## Conclusion

This proof-of-concept study demonstrates the effectiveness of a marked point process (MPP) representation of local field potentials (LFP) to mitigate the effect of volume conduction. By thoroughly characterizing oscillatory bursts in terms of their structure, signal-to-noise ratio, and transiency, we minimize the impact of volume conduction, wherein volume conducted activity do not adhere to the specific statistical properties of these transient oscillations. We further validate our approach by demonstrating that the MPP representation not only preserves the behavioral and functional information encoded in the LFPs but also introduces high resolution time-specific tools within the MPP feature space. These tools facilitate novel neurophysiological interpretations while offering computational simplicity through their sparse and compact representation of the complex, high-dimensional LFP data.

## Supporting information

**Fig S1.**
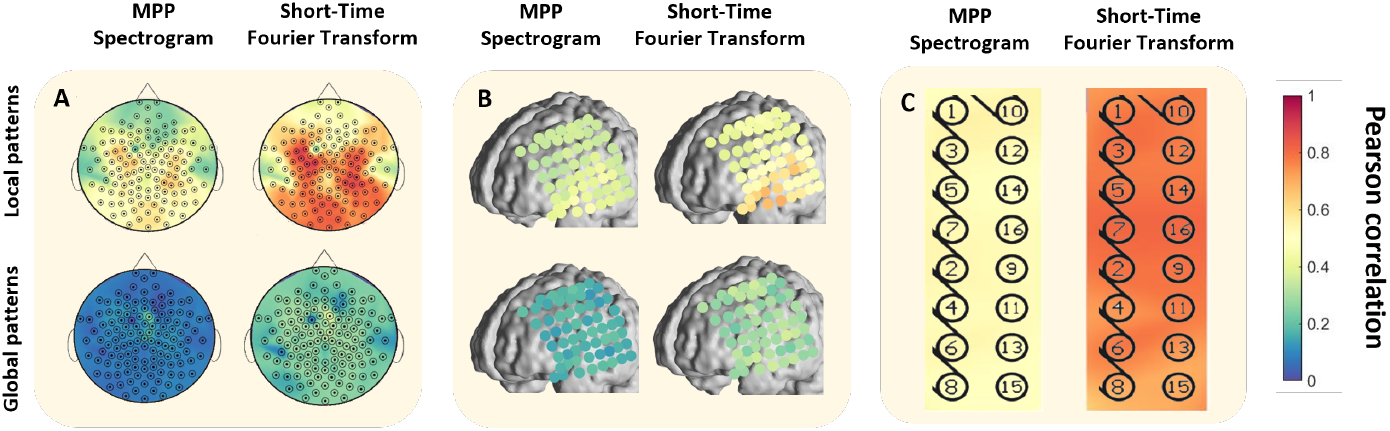
Comparison of cross-channel correlations between beta oscillations across different sensing modalities and models. **A**, Top: Local correlations—Average Pearson correlation coefficient of each EEG channel with its six nearest neighboring channels, evaluated using (left) a marked point process (MPP) representation of bursts and (right) short-time Fourier transform (STFT). Bottom: Global correlations—Average Pearson correlation coefficient of each EEG channel with its 20 farthest channels, evaluated using (left) MPP representation and (right) STFT.**B** The same analysis as in **A** applied to ECoG data from the left fronto-parietal area in a human, using the 6 nearest neighboring channels to evaluate local correlations (Top) and the 10 farthest channels to evaluate global correlations (Bottom). (**C**) The same analysis as in **A** applied to LFPs recorded in the prefrontal cortex of a rat, using the 6 nearest neighboring channels to evaluate local correlations. Data shown is from Subject 1 in each dataset.

**Fig S2.**
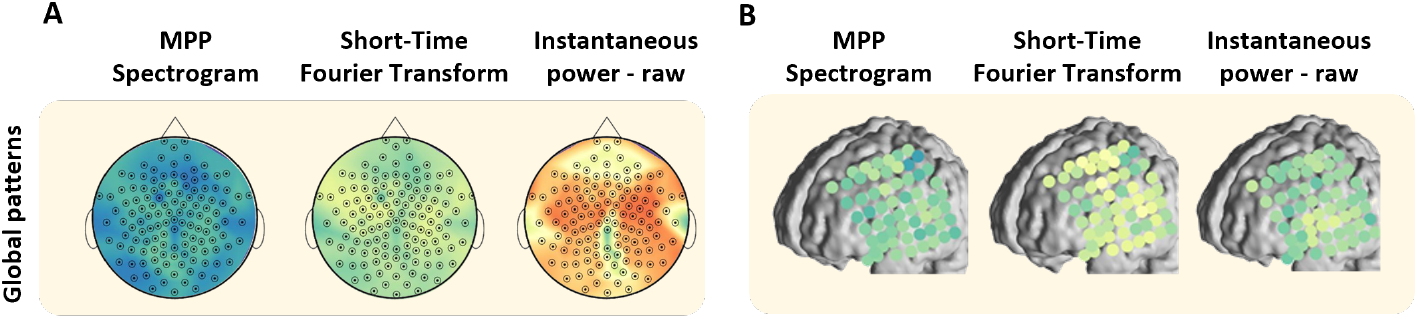
Comparison of global cross-channel correlations between high-gamma oscillations across different sensing modalities and models. **A**, Average Pearson correlation coefficient of each EEG channel with its 20 farthest channels, evaluated using (left) a marked point process (MPP) representation of bursts, (middle) short-time Fourier transform (STFT), and (right) raw EEG for an example subject. **B** The same analysis as in **A**, applied to ECoG data from the left fronto-parietal area in a human, using 10 farthest channels, respectively. Data shown is from Subject 1 in each dataset.

**Fig S3.**
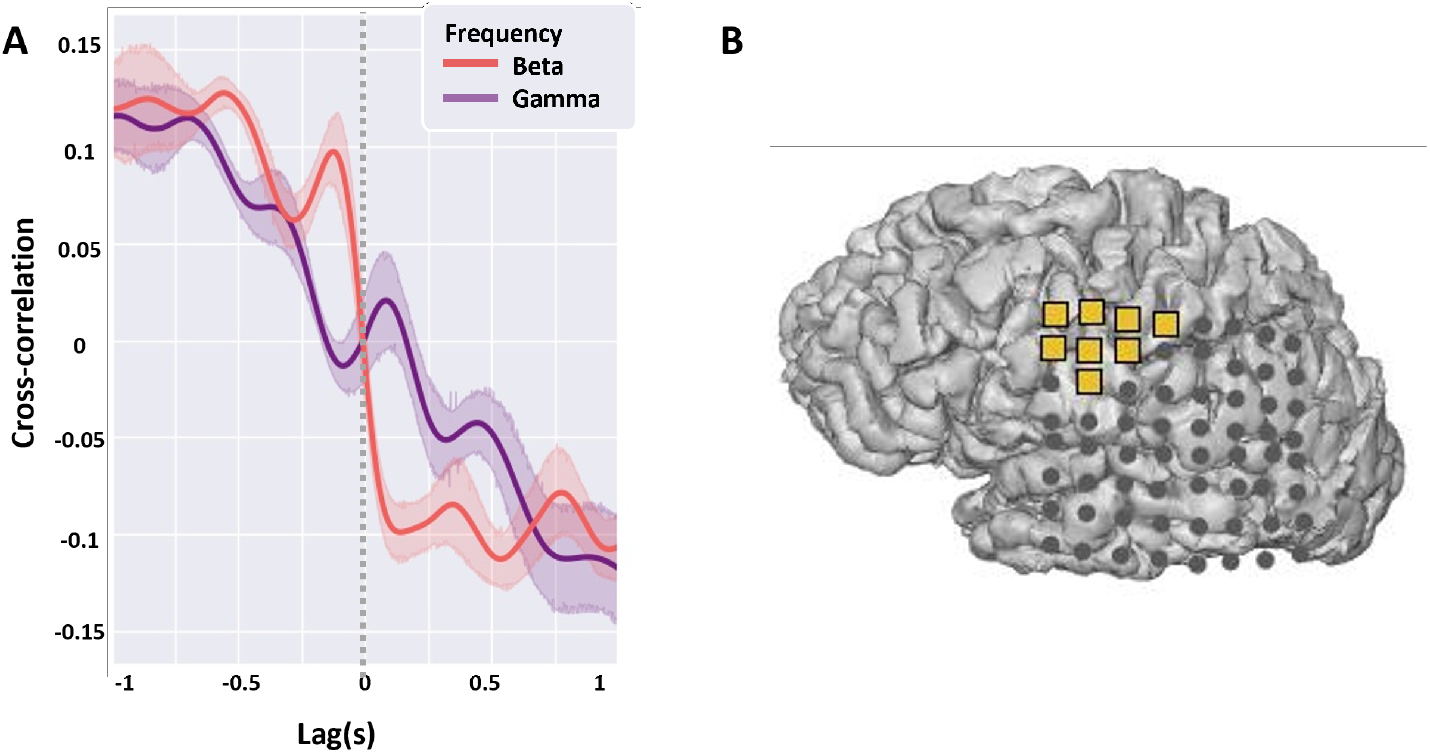
Dataset-2: Exclusion of data from subjects 4, 5 and 6: **A** Poor gamma-beta entrainment does not allow for identification of movement start time in Subject 4, resulting in poor movement classification that further compounded the errors in finger-label classification **B** In subject 5, finger-label classification was not possible due to fewer channels covering the motor cortex (yellow markers). Trials from Subjects 4 and 5 were included in all analyses except for the online classification performance. Finally, because Subject 6 had only 8 trials per class, it was excluded from all analyses, as the limited sample size would not yield statistically significant results.

**Table S1.** Subject and experiment overview of the ECoG dataset (Dataset-2, Fig 1**C**)

| ID | Age | Gender | Handed-ness | Hemisphere coverage | No. of electrodes | Trials per class |
| --- | --- | --- | --- | --- | --- | --- |
| 1 | 18 | Female | Right | Left - Fronto-parietal | 46 | 30 |
| 2 | 21 | Male | Right | Right - Fronto-temporal | 63 | 30 |
| 3 | 27 | Female | Right | Left - Fonto-temporal-parietal | 61 | 30 |
| 4 | 35 | Female | Right | Left - Fronto-temporal | 58 | 22 |
| 5 | 26 | Male | Right | Left - Parietal-temporal-occipital | 64 | 30 |
| 6 | 45 | Female | Right | Left - Fronto-temporal | 43 | 8 |
| 7 | 32 | Male | Right | Left - Fronto-temporal-parietal | 64 | 30 |
| 8 | 19 | Female | Right | Right - Fronto-parietal | 38 | 17 |
| 9 | 18 | Female | Right | Left - Frontal | 47 | 26 |

**Table S2.** Decoder accuracy for finger classification in the channel and the corresponding brain area with the highest accuracy.

| ID | Accuracy | Channel Location |
| --- | --- | --- |
| 1 | 0.36 | Dorsal M1 |
| 2 | 0.45 | Dorsal M1 |
| 3 | 0.45 | Dorsal M1 |
| 4 | 0.40 | Dorsal M1 |
| 5 | 0.40 | Dorsal S1 |
| 7 | 0.47 | Dorsal M1 |
| 8 | 0.54 | Dorsal M1 |
| 9 | 0.48 | Frontal (non-rolandic) |

## Acknowledgments

This project was primarily funded by the National Science Foundation (grant number: 1631759 to J.C.P). We thank Karim Oweiss and Andreas Keil for their insightful discussions, as well as Ali Mohebi and Kierstin Riels for their generous data sharing.

